# Theoretical and numerical investigation of the consistency of model comparisons in pharmacometrics

**DOI:** 10.1101/2025.10.18.683202

**Authors:** Lena M. Appel, Bernhard Steiert

## Abstract

Pharmacokinetic (PK) and pharmacodynamic (PD) models are essential tools in drug development, making the selection of an appropriate model critically important. When using likelihood ratio tests (LRTs) to compare nested models, it is crucial to ensure their validity, especially when parameters are fixed. This work examines the continuity of likelihood functions as a necessary condition for LRT validity within the framework of population modeling. By decomposing the Objective Function Value (OFV), we identify scenarios where parameter fixing leads to non-continuous likelihood behavior, potentially invalidating the LRT application. A proof and numerical examples illustrate that while fixing population parameters maintains continuity through compensatory behavior of terms within the OFV, fixing individual parameters introduces discontinuities. Overall, this work underscores the need for careful consideration of parameter fixation in population models: It shows that population parameters can be fixed without violating the continuity condition for LRTs and suggests that introducing covariates may provide a viable alternative for fixing individual parameters. Further investigation into the sufficiency of continuity as a condition for the LRT’s validity is needed.

## 1 Introduction

Pharmacokinetic (PK) and pharmacodynamic (PD) models are essential tools in drug development, guiding critical decisions in pharmaceutical research and clinical practice [6, 2]. Selecting the most appropriate model from numerous possibilities is a crucial step. The likelihood function serves as a primary metric for evaluating different model variants and optimizing parameters to best describe observed data.

As standard of practice, pharmacometricians employ statistical tests, to compare models and identify the most suitable one for simulations and predictions.

When comparing nested models - where one model is a simplified version of another - the likelihood ratio test (LRT) is a widely employed statistical method. E.g. a model with fixed parameters is considered “nested” within a more general model where those parameters are estimated. It is important to note that in classical regression analysis, fixing a parameter typically reduces model flexibility, leading to an equal or worse likelihood. The LRT, calculated as the difference in log-likelihoods between nested models, is a powerful tool for the comparison of nested models [7]. However, the validity of the LRT, particularly when parameters are fixed, is not always guaranteed within the complex framework of population modeling.

A fundamental condition for the validity of the LRT is the *continuity* of the likelihood function with respect to its parameters. This means that infinitesimal changes in parameter values should only result in slight deviations in model predictions and, consequently, minimal changes in the likelihood and the LRT statistic. For inherently continuous functions, like ordinary differential equation (ODE) models, implicit dependencies of the likelihood on parameters generally maintain this continuity. Therefore, in a classical non-population approach, the continuity requirement for a valid LRT is typically met.

However, the context shifts significantly in pharmaceutical research with the widespread use of the *population approach*, also known as nonlinear mixed-effects (NLME) modeling. This approach accounts for inter-individual variability by assuming that individual parameters follow a distribution around a population typical value. While powerful, this hierarchical structure adds substantial complexity to the likelihood function, making its exact computation challenging and computationally expensive[8, 5, 9]. Consequently, approximations such as the First-Order (FO) or the First-Order Conditional Estimation (FOCE) method, often implemented in software like NONMEM^®^, are commonly employed. In this work, we will focus on these approximations. Unlike classical regression, these approximated likelihoods, from which the Objective Function Value (OFV) is calculated, contain additional terms which quantify the distribution of the individual parameters [11, 1]. These include explicit dependencies on parameters and can therefore lead to the violation of *continuity*, potentially invalidating the LRT as a reliable test statistic for model comparison.

### Motivation

Despite the critical role of LRTs in population modeling for model selection, previous research has not comprehensively investigated the impact of the population approach’s complex likelihood on LRT validity. More precisely, the specific implications of fixing parameters within this framework remain not well understood. This knowledge gap arises partly from the lack of available details about the composition of the approximated likelihood, and partly because the non-continuous behavior of the likelihood only shows when individual parameters are fixed. Such scenarios, while important for specific applications like sequential analysis for population PK/PD [12], are rarely thoroughly investigated. Nevertheless, a good understanding of the underlying mathematical equations and their behavior is crucial to ensure reliability of model comparisons in pharmacometric research.

To address this critical gap, the objectives of this work are:

1. To decompose the likelihood function of the FOCE approximation for NLME models.
2. To examine individual likelihood components for non-continuous behavior that could compromise the validity of LRTs.
3. To provide guidance on the scenarios under which LRTs remain valid within the population modeling framework.

It is important to note that this manuscript primarily investigates *continuity* as a necessary condition for a valid likelihood ratio test statistic; other necessary or sufficient conditions are beyond the scope of this analysis.

## 2 Methods and assumptions

### 2.1 Derivation of OFV and its decomposition

To understand whether discontinuities could emerge when removing or fixing a parameter, we decompose the NLME likelihood obtained under the first order conditional estimation (FOCE) approximation. A derivation of the approximated likelihood in NONMEM can be found in [11], and further information on how these components can be obtained and analyzed in NONMEM can be found in [1].

### 2.2 Assumptions and definitions

When fitting an PK/PD model, like an ODE model, to a data set, the underlying assumption is that the data follows a certain probability distribution and a statistical model is built, including parameters to describe the distribution. Then, the probability to obtain the data, *p*(*data*|*parameters*), can be formulated. The likelihood *L*(*parameters* |*data*) is obtained, when the probability density *p* is interpreted as a function of the parameters. When a population approach is applied, there are two types of parameters: It is assumed that the parameters describing each individual are following a distribution around a corresponding population parameter.

The likelihood function *L*(Θ, Σ, Ω), which is discussed in more detail in the next section, is then described by the following components:

- Θ: fixed effect parameter, typical values for the population
- Σ: residual variance-covariance matrix (Square matrix with dimensions of observed data as the variance corresponds to the uncertainty of the observed data)
- *η*: random effect parameter, representing the deviation of an individual subject’s parameter from the typical value of the population. Importantly the individual parameter value *P*_*i*_ is constructed from *η* and the population typical value Θ_*T V*_ . The most common assumption is that the individual values are log normally distributed leading to the following relationship *P*_*i*_ = Θ_*T V*_ *·* exp (*η*_*i*_).
- *y*: observed data vector
- *f* : predicted function value (IPRED) corresponding to the observed data *y*
- Ω: the variance-covariance matrix for the interindividual variability.

In the following, *f* is assumed to be twice continuously differentiable with respect to the parameters. Furthermore, Σ is assumed to be finite valued, i.e. there is a finite error on the data the prediction function is fitted to. Both assumptions are typically valid for NLME pharmacometric models.

If *f* is twice continuously differentiable with respect to the parameters, then the likelihood is also twice continuously differentiable with respect to the parameters for ordinary (non-population) regression analysis. However, this changes for the population approach, even if *f* is twice continuously differentiable, as there is an explicit dependency of the likelihood on the parameters. Terms with explicit dependencies give rise to the critical terms and are analyzed.

### 2.3 Critical cases which can result in discontinuity of OFV

After identifying critical terms within the OFV which can cause discontinuous behaviour, the behaviour of these terms is examined when fixing parameters to different values.

More precisely, first, the case of a component of Ω approaching zero is analyzed. Then, by setting Ω to zero the model is reduced and a nested model is obtained and the behavior of the likelihood is tested. Second, the case is considered where individual parameters are manually specified for each individual, i.e. *P*_*i*_s are specified, instead of estimating them. The manual specification can be implemented by providing individual values for a parameter (e.g. clearance) in a column of the data set, which are different for each individual *i*, fixing the parameter in the script (*P*_*CL,i*_) to the values in the column and then fitting the remaining parameters. This column is comparable to the data column provided when introducing individual covariates. A special case would be to use the Empirical Bayes Estimates (EBEs) from a previous estimation run and fixing *P*_*x,i*_s to these values (*x* denoting a parameter like clearance). The corresponding Ω_*x*_ entry becomes obsolete and is removed from the Ω matrix. This procedure is depicted in Figure 1 and will be termed as “fixing *P*_*i*_s to covariates” from now. The influence of this procedure on the likelihood is analyzed to assess the validity of the LRT.

**Figure 1.**
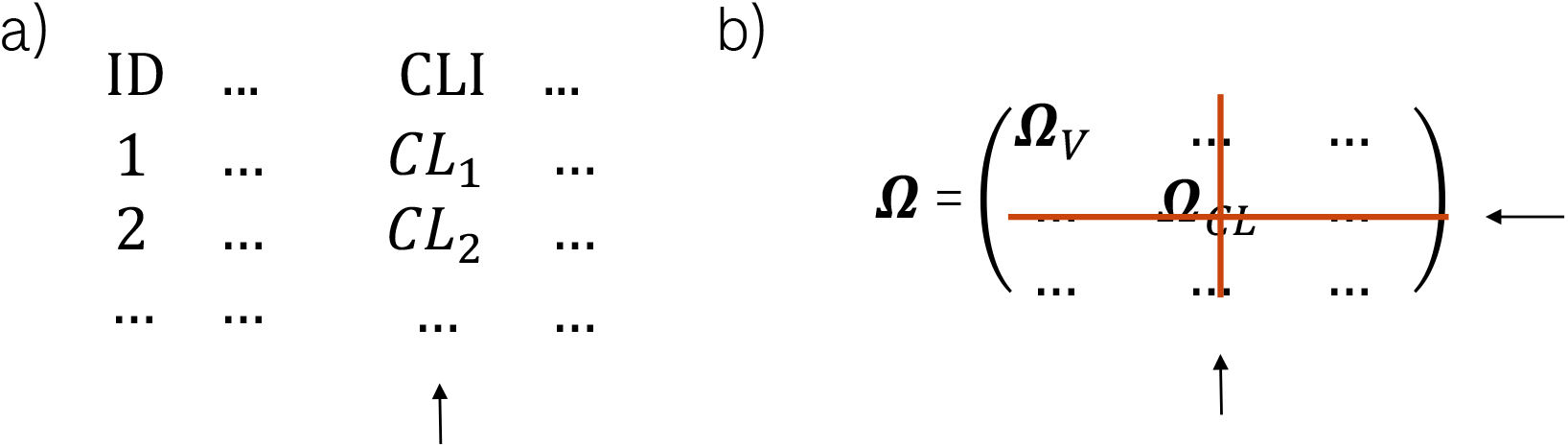
a) In the data file a column is inserted with individual values for a parameter (here: Clearance) b) The corresponding Ω value (here Ω_*CL*_) in the covariance matrix becomes obsolete as neither the population parameter for the clearance nor its interindividual variability is estimated

### 2.4 Example to confirm theoretical considerations

Finally, an example of a pharmacokinetic/pharmacodynamic (PK/PD) model is used to illustrate the changes in the OFV components in various scenarios. The PK part is described by a two compartment model with linear absorption and the PD part consists of an indirect response model. The analysis is conducted with the FOCE approximation with interaction, as a more general case of FOCE without interaction for which the theoretical calculations have been conducted.

In a first step the PK/PD model is used to generate a simulated data set with 100 individuals. A single simulation suffices to investigate the general dependency of the OFV on changes of the parameters. In a second step the generated data is fitted under different conditions.

The PK part of the model has no interindividual variability and is fixed throughout the whole simulation and fitting (sequential model). The PD part of the model constitutes an indirect response model using 1) the drug concentrations from the PK model, 2) population parameters, and 3) interindividual variability. For the simulation, the parameters of the indirect response model are chosen such that the dose response relationship is sensitive to when parameters are changed, see supplementary section B.

When fitting the following scenarios are evaluated to test for the occurence of non-continuous behaviour of the OFV when changing from the baseline scenario (first scenario):

Sc1 all parameters are estimated

Sc2 one Ω or Θ is fixed to the estimated value from Sc1

Sc3 one Ω is fixed to various smaller values

Sc4 one Ω is fixed to zero or the corresponding interindividual variability is removed entirely by removing this Ω

Sc5 in addition to the removal of an Ω the corresponding individual parameters (*P*_*i*_s) are set to the estimated values from Sc1 which is called “fixing *P*_*i*_s to covariates”, see 2.3.

The fitting is conducted with NONMEM^®^ v7.4.3 and for the analysis of the NONMEM output and simulations, R Statistical Software v4.1.2, see [10], is used. To read the output from NONMEM and calculate the contributions of the different terms of the OFV, a script published in [1] has been adapted.

## 3 Results

### 3.1 Detailed composition of the likelihood function and OFV

#### 3.1.1 The NLME likelihood function

The NLME modeling approach (population approach) uses a hierarchical model due to the assumption that the individual parameters follow a distribution around a population typical value. In particular, both the parameters and the data are assumed to be sampled from an underlying distribution such that the joint population likelihood function

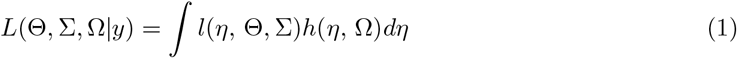

contains the product of two partial likelihoods *l* (*η*, Θ, Σ) and *h*(*η*, Ω).

The partial likelihood *l*(*η*, Θ, Σ) is defined by the conditional probability density *p*(*y* |*η* Θ Σ) of the data *y* given the model *f* = *f* (*η*, Θ, Σ) with the random effect parameters *η*, the population typical parameters Θ and the residual variance-covariance matrix Σ. *p*(*y* |*η*, Θ, Σ), which is a function of the data *y* can be interpreted as a function of the parameters leading to the likelihood function *l*(*η*, Θ, Σ|*y*). Assuming a Gaussian error distribution we can write *l*(*η*, Θ, Σ|*y*), which accounts for the distance between the evaluated function values and the data, as

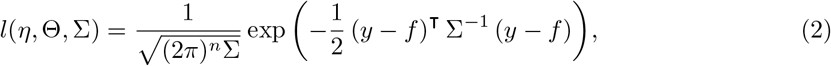

with *n* the number of observations.

The second partial likelihood stems from the probability density of the individual parameters *η* which are assumed to be normally distributed with a variance Ω:

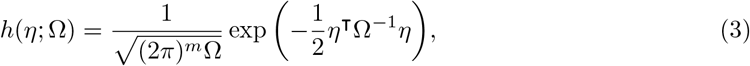

with *m* the number of parameters.

#### 3.1.2 Approximation of the NLME likelihood and presentation of its components

As the exact computation of this integral (1) is difficult, NONMEM uses the Laplacian approximation of the integral, of which FOCE is a further approximation, see also [11]. In this work we will focus on this approximation and an additional error model.

First, the Laplacian approximation and two times the negative logarithm is applied to the likelihood *L*(Θ, Σ, Ω). Then, after omitting constant terms which do not contribute to the minimization of the expression, the expression for the likelihood from which the OFV is calculated, is obtained as

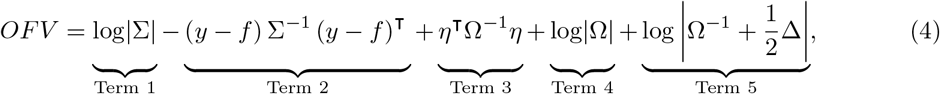

containing Terms 1 - 5 that are examined individually in the following. Δ is the Hessian matrix of the partial likelihood *l*(*η* Θ Σ).

Note that the frequently used FOCE algorithm is a further approximation of the equation (4) where the Hessian matrix Δ is approximated by a function of the gradient vector,

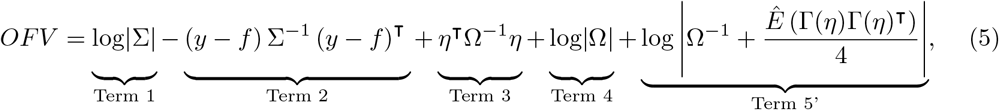

see [11]. Here, *Ê* (Γ(*η*)Γ(*η*)^T^) is the expectation taken relative to *y* and *η* and Γ(*η*) is the gradient of −2 log *l*(*η* Θ Σ).

#### 3.1.3 The interpretation of the OFV components

Generally speaking the aim is to choose parameters such that the sum of terms 1 - 5 in the OFV is minimized. Given a set of parameters, changing these can lead to an increase or decrease of the value of these terms, where an overall increase would be “penalizing” the choice of parameters. An overview in which cases the different terms decrease is given in Tab. 3.1.3. It becomes obvious that some terms change in exactly the opposite directions when parameters are changed in a certain way such that a compromise needs to be found.

Term 1 and term 2, which are stemming from *l*(*η* Θ Σ), are terms which appear in the negative loglikelihood of non-population approach regressions, as well. Term 2 is penalizing a choice of parameters which increases the distance of the model prediction to the actual measured data points for all individuals. The term is divided by the parameter for the residual error, favouring also a large residual error, Σ. While Term 2 benefits from a large residual error, Term 1 is penalizing the choice of a large Σ, thus forcing the optimization to find a compromise and preventing the choice of an arbitrarily large error.

In addition to term 1 and 2, in a population approach it is assumed that the individual parameters *P* follow a distribution around the population parameter Θ. This is realized by constructing *P* from *η* which is distributed with mean zero and variance Ω. Thus, term 3 and term 4 resemble term 2 and term 1, respectively, in so far as term 3 leads to a minimization of the distance between *η* and zero (and by this, the distance between individual parameters and the population parameter) and favor a large Ω by which it is divided, while the choice of a too large interindividual variability Ω is penalized by term 4.

As a part of the Laplace approximation the second derivative of log (*l*(*η* Θ Σ)*h*(*η* Ω)) is calculated and enters as term 5.

### 3.2 Non-continuous behavior can be expected due to explicit parameter dependencies

In the introduction the importance of the continuity of the model and the likelihood function with respect to the parameters is explained in order for the likelihood ratio test (LRT) to be *continuous* and therefore a valid metric. This requirement is typically met for commonly applied PK/PD models, see also 2.2. In a non-population approach regression analysis the likelihood, which has implicit dependencies on the parameters, would then also be continuous and the necessary condition for the LRT to be a valid test statistic would be met.

**Table 1:**
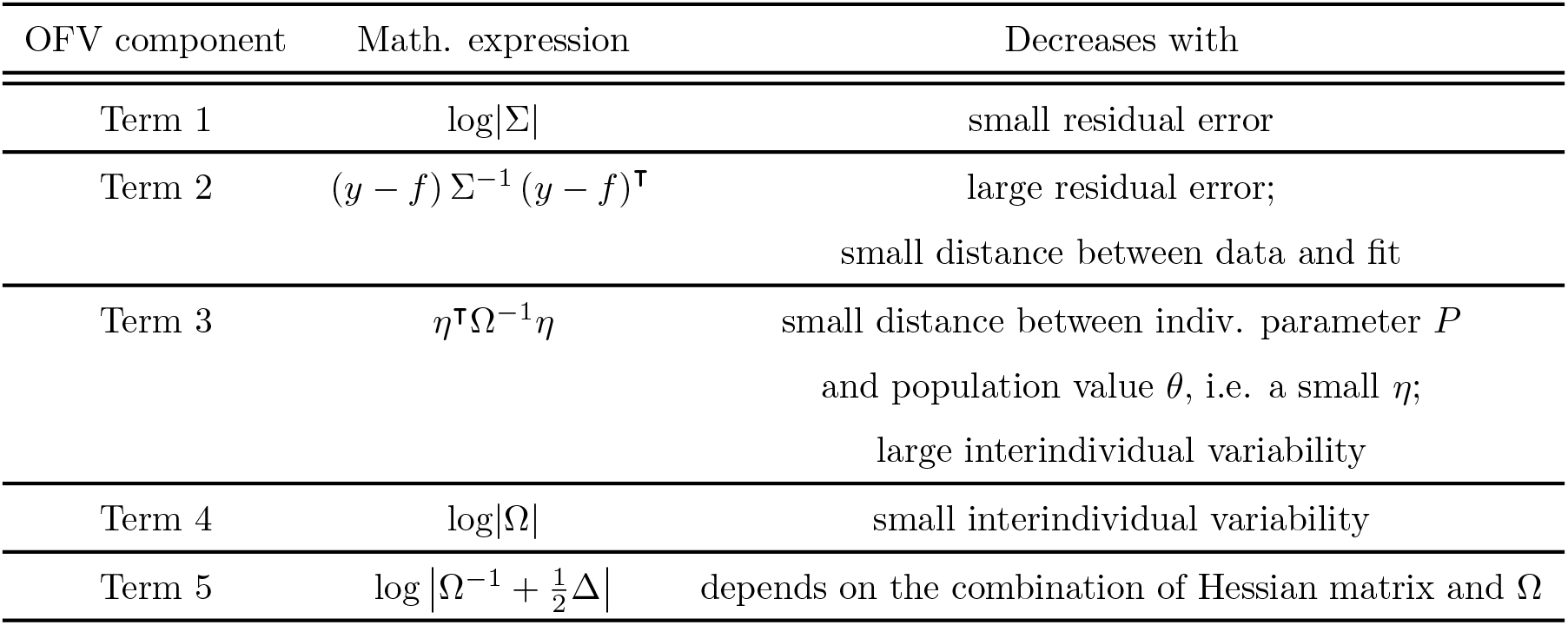
Summary of the influence of different OFV components to the overall OFV, which needs to be minimized during a fit. It is outlined how the choice of parameters influences their contribution to the OFV.

However, in population approach, the likelihood function has <u>implicit</u> and <u>explicit</u> dependencies on population parameters, see eq. (5). In contrast to implicit dependencies, explicit parameter dependencies can give rise to an abrupt change of the likelihood, e.g. it can show divergent behavior as the parameters approach certain values, leading ultimately to singularities when parameters take on this value. This could lead to the likelihood to be non-continuous and, thus, disqualify the likelihood ratio to be used as a test statistic in these cases.

From this analysis, we conclude that there is a need to identify and further examine cases in which terms in the OFV with explicit parameter dependencies could give rise to non-continuous behavior.

### 3.3 Non-continuous behavior can be expected when fixing parameters

Generally, fixing a parameter and subsequently refitting the remaining parameters influences the squared distance of the model fit from the data, as a degree of freedom is removed. Hence, the model fits either the data equally well, if the parameter is fixed to the maximum likelihood estimated value (or if the parameter is fixed to another value, but the parameter is non-identifiable) or the model fits more poorly when the parameter is suboptimal. But the OFV should not improve. Furthermore, as the parameter moves away from the optimal value, the gradual worsening of the fit should be reflected by continuous increase of the OFV.

But this is not always the case in a population approach and introduces another case where the LRT looses its validity, as explained in the following.

### 3.4 The OFV terms explicitly depending on Ω could give rise to singularities and discontinuous behavior, but only when fixing individual parameters

In the following it is detailed why an explicit dependency on parameters can lead to singularities (and thus, a discontinuity of the OFV) and which terms of the OFV and cases of fixing parameters need further investigation.

#### 3.4.1 Term 1 and term 2 without an explicit dependency on critical parameters do not give rise to singularities

As stated in 2.2 it can be assumed that for a pharmacometric model, f is continuous with respect to its parameters and that Σ is well-behaved, i.e. not taking on extreme values including zero. In contrast, Θ could be zero (when a reaction does not take place), Ω could be zero when there is no IIV for a parameter, or *η* could be zero when there is no data for an individual to justify a non-zero value, i.e. shrinkage is high for this individual.

As there is no explicit dependency from Θ, Ω or *η* in the term 1 and term 2 in eq. (4), we conclude there will be no divergence or singularity leading to a non-continuous behavior as the parameters change. In an ordinary regression without a population approach there are only term

1 and term 2, which is why the problem of a non-continuous behaviour does not exist and has therefore not been investigated in the literature.

#### 3.4.2 Ω approaching zero could in principle give rise to singularities in term 4 and term 5

The explicit dependency on Ω of Terms 3, 4, and 5 in eq. (4) can lead to a divergence as Ω→0 and a singularity at Ω = 0, because Ω is in the denominator or the logarithm of Ω is taken. Term 3 however is independent of Ω since *η* ∼ *N* (0, Ω) such that *η*^T^Ω^−1^*η* is merely a normalization of the parameter *η*. To this end only Term 4 and Term 5 in eq. (4) need to be regarded as Ω → 0. However, it is shown in section 3.5 that the divergence of the last two components are compensating each other.

#### 3.4.3 Changing the structure of Ω as a consequence of fixing individual parameters can give rise to a non-continuous change in terms 3, 4 and 5

The explicit dependency on Ω gives rise to another unexpected non-continuous behavior. It also violates our expectation that the OFV should not improve if parameters are fixed to previously estimated values, while the remaining parameters are refitted, as stated in section 3.3.

More precisely, the unexpected behavior occurs due to terms 3, 4, and 5 when we “fix *P* s to covariates”: In a first run Ω is estimated to a value Ω_*est*_ with corresponding *η*_*est*_ and resulting individual parameters *P*_*est*_. In the next fit, the individual parameters, *P* s, are fixed to the *P*_*est*_ while other parameters are refitted. Normally, it is expected that fixing a parameter to an estimated value and refitting leads to the same likelihood because the squared distance of the model from the data - and the ability of the model to describe the data - should not change.

As expected, in this procedure, the squared distance of the model to the data, described by terms 1 and 2 which constitute the non-population likelihood, stay the same. But the corresponding entry in Ω is omitted, as it becomes obsolete, and terms 3, 4 and 5 change abruptly due to their explicit dependence on Ω. And together this leads to a non-continuous change in the overall OFV - frequently even improving the OFV - and invalidates the LRT. A corollary of the proof which primarily examines the behavior of the OFV for Ω *→*0 in section 3.5 explains this non-continuous behavior.

### 3.5 Proof: the divergent behavior of term 4 and term 5 in the OFV compensate as Ω approaches zero, but fixing individual parameters leads to a discontinuous behavior

In the following it is proven that the transition of an entry of Ω approaching zero and being omitted is continuous, i.e. that diverging terms are compensating in the OFV, see eq. (4). The focus of the proof therefore lies on the critical terms discussed in 3.4 which have an explicit dependency on Ω, i.e. Term 4 and Term 5 in eq. (4).

More precisely, here it is shown for a diagonal Ω matrix that as an entry of Ω approaches zero and leads to a divergent behavior, the corresponding terms cancel out. Since all contributions from this entry in Ω cancel out as it approaches zero, there is a smooth transition to the OFV value where the entry in Ω is being omitted entirely.

A corollary of this proof also explains why replacing a finite-valued entry of Ω by “fixing *P*_*i*_s to covariates” would lead to a non-continuous change in the OFV and hence an invalid LRT.

Note that the proof in the following works analogously for (5), where the Hessian matrix is replaced by the expectation value of the gradients.

An overview of key logarithmic and determinant calculation techniques applied in this work can be found in the appendix, see A.

#### 3.5.1 Proof for continuity of OFV values as a component of Ω approaches zero

Without loss of generality it can be assumed that the entry of the diagonal matrix Ω approaching 0 asymptotically is the first entry and the entry is termed *a*.

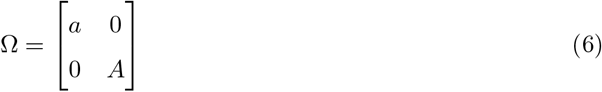

with *A* being a diagonal matrix with all other Ω entries.

We will now prove the equivalence of OFV in 3 scenarios: *a* approaching zero, *a* being set to zero, and omitting the corresponding entry in Ω, i.e.

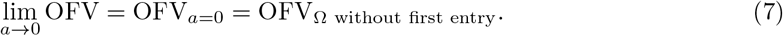

The transition between *a→* 0 and *a* = 0 is expected to be smooth for Terms 1 - 3 as discussed in 3.4, such that the focus lies on the sum of Term 4 and Term 5, which may become singular for *a* = 0. We define the sum as

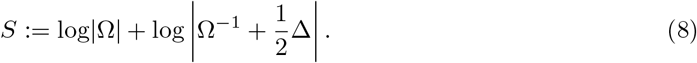

Further define

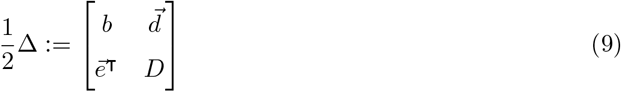

where 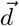 and 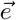 are the first row and first column of 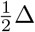. Then, S can be written as

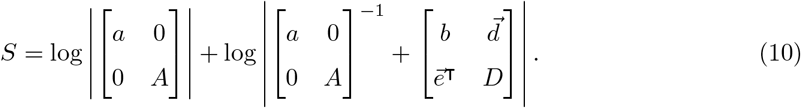

To understand the implications of omitting the first column and row of Ω including *a*, a closer look at Δ is needed. It contains the second derivatives with respect to the *η*:

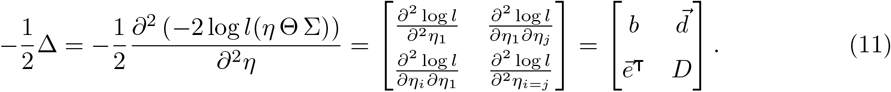

Now, *η* is distributed as *η ∼ N* (0, Ω) and *η*_1_ *∼ N* (0, *a*). Therefore, all entries in the first row and column of Δ, namely *b*, 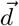 and 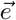 are second derivatives with respect to *η*_1_ and disappear, if the first entry of Ω, *a*, is taken out.

Focusing on terms 4 and 5, that is S, what remains to be shown from eq. (7) is:

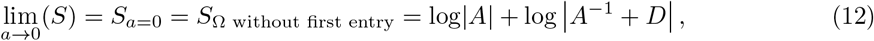

which is proven in the following.

**Proof** Using the fact that the inverse of the diagonal matrix is the diagonal matrix of the inverted entries, eq. (8) can be written as:

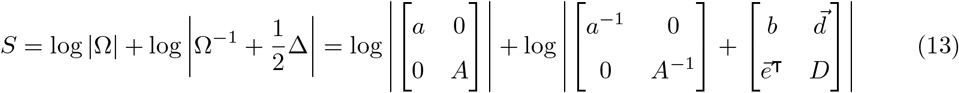

and simplified as

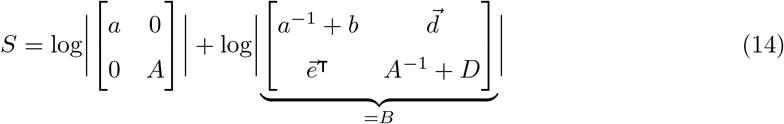

and using logarithm and determinant rules the following equation is obtained:

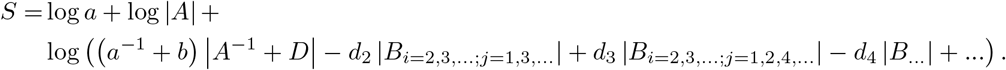

Here, the matrices *B*_*i*=2,3,…;*j*=1,3,…_ and *B*_*i*=2,3,…;*j*=1,2,4,…_ are submatrices of *B* and do not contain the first row and the xth column of *B* and are multiplied with the corresponding entry *d*_*x*_ of the vector 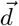 respectively. *B*_*i*=2,3,…;*j*=1,3,…_ and *B*_*i*=2,3,…;*j*=1,2,4,…_ will be abbreviated with *B*_1;2_ := *B*_*i*=2,3,…;*j*=1,3,…_ and *B*_1;3_ := *B*_*i*=2,3,…;*j*=1,2,4,…_ in the following, where the missing rows and columns are indicated by the subscript. Most importantly, the submatrices of *B* do not contain *a*. The expression can further be rewritten as:

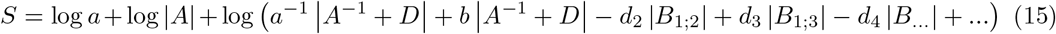

and the last expression in logarithm brackets is divided by *a*^−1^ and using the logarithm rules *a*^−1^ is taken out of the log bracket. This returns

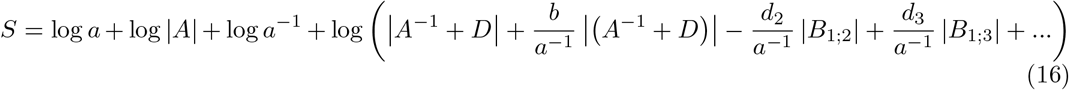

and log *a* and log *a*^−1^ cancel out, such that

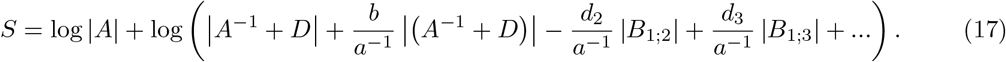

This expression can be studied for the case of *a →* 0, that is *a*^−1^ *→ ∞*. Expressions with *a*^−1^ in the denominator will approach zero as *a →* 0, returning

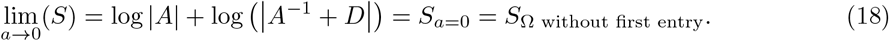

This is the desired result, see eq. (12).

Thus, it can be seen that

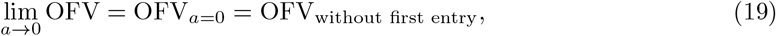

what was needed to be proven.

#### 3.5.2 Corollary: Fixing *P* to covariates does not lead to a smooth transition of the OFV

As stated in 3.4.3, when “fixing *P* s to covariates” the corresponding interindividual variability is omitted, that is the Ω entry is taken out, which would be “*a*” in the proof above. In eq. (17) it can be seen that taking out “*a*” without its approaching zero leads to an abrupt change in the OFV, as the terms with 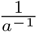 in eq. 17 are not approaching 0 and are therefore unequal to *S* or OFV_without first entry_.

That is, for *a* ≠ 0

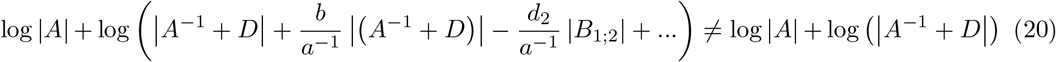

and thus,

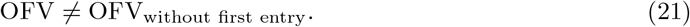

## 4 Numerical example

We now confirm the findings and theoretical calculations by looking at the results of a numerical example: To this end, the following scenarios with the model described in the Suppl. B are analyzed:

Sc1 all parameters are estimated

Sc2 Ω_*i*_ and a Θ_*i*_ are fixed to the estimated value, but the other parameters *j* ≠ *i* are fitted

Sc3Ω _*i*_ is fixed to decreasing smaller values while other parameters *j* ≠ *i* are fitted

Sc4 Ω _*i*_ is fixed to zero or removed, i.e. the corresponding interindividual variability is removed

Sc5 Ω_*i*_ is removed, but the interindividual variability is re-introduced by setting the corresponding individual parameters (*P*_*i*_s) to the estimated values (EBEs) from Sc1 - a process which is termed “fixing *P*_*i*_s to covariates”, see 2.3. To obtain a similar fit as Sc1, population parameters are mostly fixed to values estimated in Sc1.

The Ω entry which is varied or set to zero in this numerical example is responsible for the interindividual variability of EC50. A summary of the behavior of all OFV component values for the different scenarios can be seen in Suppl. Fig. 4. All scripts used to calculate the scenarios can be found in Suppl. C.

We observe the following results.

### Result 1: the presented OFV decomposition is confirmed

The numerical example confirms that the sum of the five OFV components calculated as described in Eq. 4 with a script adapted from [1], is equal to the OFV value returned by NONMEM.

### Result 2: fixing a population parameter to its estimated value has no influence on the OFV (Sc2)

Fixing population parameters Ω_*EC*50_ and Θ_*EC*50_ to the estimated values from Sc1 results in the same OFV values. This is confirmed for both the overall OFV, see dashed lines in the two middle panels in Fig.2 (Sc1 and Sc2), and for the individual OFV components: For term 4, term 5 and term 4+5 see dashed lines in the two middle panels in Fig.3 (Sc1 and Sc2) and the behavior of all other OFV components can be seen in Suppl. Fig.4.

### Result 3: the likelihood changes continuously when fixing Ω to decreasing finite values

When fixing an Ω_*EC*50_ to steadily decreasing smaller values (Sc3), the components 4 and 5 diverge in opposite directions, but these divergent behaviors cancel out when added, see second panel in Fig. 3 (Sc3). Thus, the overall OFV does not diverge.

### Result 4: the likelihood changes continuously when fixing Ω to extreme values, including zero

When comparing the OFV components in the limit of Ω_*EC*50_ approaching zero (Sc3) and Ω_*EC*50_ being set to zero (Sc4) the following is observed: The term 4 and 5 diverge and the change when set to 0 is sudden, but the sum changes continuously. This can be seen by comparing the first two panels in Fig. 3: In the first panel Ω_*EC*50_ is fixed to 0 or omitted (Sc4) and in the second panel Ω_*EC*50_ is set 10^−11^ (Sc3). Term 4 and 5 change abruptly, but the sum stays the same, see the dotted line. All other OFV components stay the same, see dotted lines in first two panels (Sc3 and Sc4) in Suppl. Fig 4. Both observations are in line with the calculation in 3.5.1.

### Result 5: fixing *P*_*i*_s to covariates leads to a non-continuous change of the likelihood

If Ω_*EC*50_ is removed and the interindividual variability is re-introduced by “fixing the corresponding *P*_*EC*50_s to covariates” (Sc5), as described in 2.3, a non-continuous change in the OFV can be observed. This can be seen by following the dashed line in the third and fourth panel in Fig. 2 (Sc1 and Sc5). As expected the residual error (term 1) and with this the distance of the function from the data points is unchanged (term 2), see dashed lines in the third and fourth panel in Suppl. Fig. 4 (Sc1 and Sc5), but the other components of the OFV change significantly, see third and fourth panel in Fig. 3 and Suppl. Fig. 4. This behavior confirms the theoretical consideration in 3.5.2.

**Figure 2.**
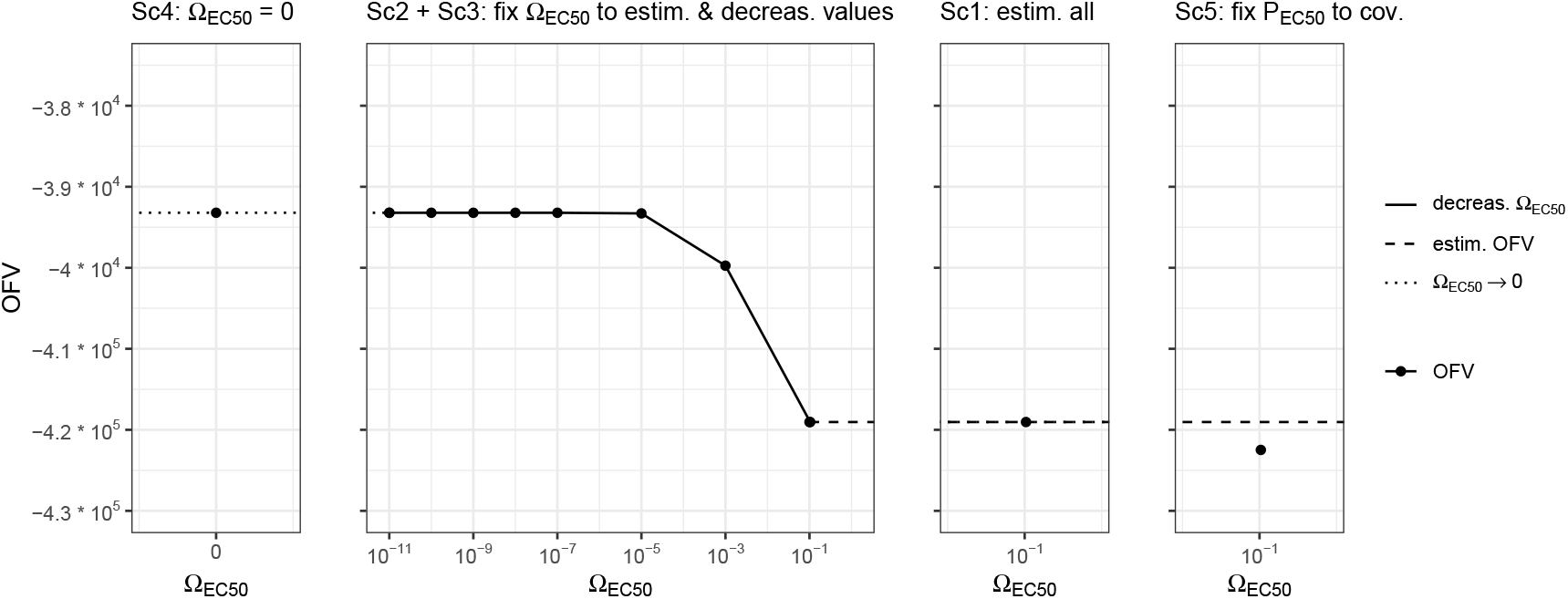
Value of OFV for different fitting scenarios Sc1-Sc5 where the Ω_*EC*50_ is estimated, fixed or omitted. **Sc1** all parameters are estimated simultaneously. **Sc2** Ω_*EC*50_ and Θ_*EC*50_ are fixed to the estimated value from Sc1 and all other parameters are refitted. **Sc3** Ω_*EC*50_ is fixed to decreasing values and a contiuous change can be seen. **Sc4** Both setting Ω_*EC*50_ to zero or omitting Ω_*EC*50_ entirely leads to the same values for all components (only Ω_*EC*50_ = 0 shown here). **Sc5** Ω_*EC*50_ is removed and the interindividual variability is re-introduced by “fixing the corresponding *P*_*EC*50_s to covariates”, see 2.3.

**Figure 3.**
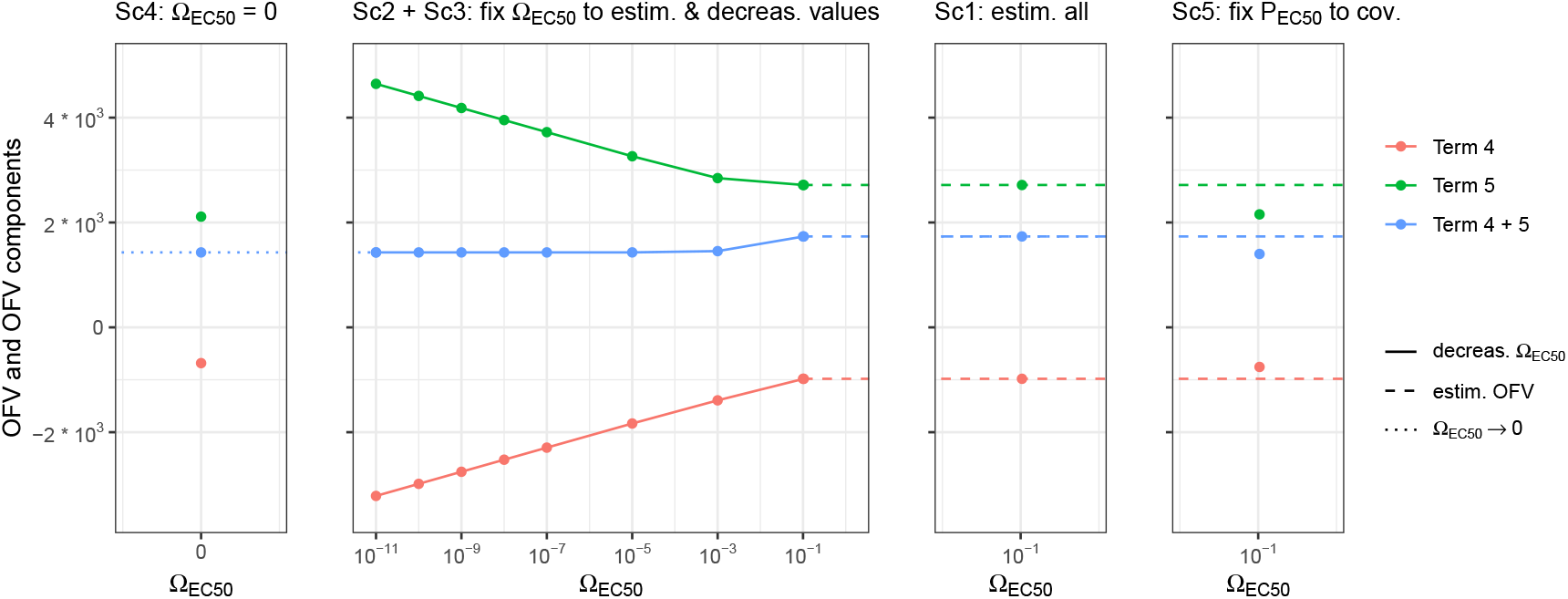
Value of term 4, term 5 and term 4+5 for different fitting scenarios Sc1-Sc5 where the Ω_*EC*50_ is estimated, fixed or omitted. **Sc1** all parameters are estimated simultaneously. **Sc2** Ω_*EC*50_ and Θ_*EC*50_ are fixed to the estimated value from Sc1 and all other parameters are refitted. **Sc3** Ω_*EC*50_ is fixed to decreasing values and a contiuous change of components can be seen. A divergence of term 4 and 5 can be seen, but the change of term 4 + 5 remains small. **Sc4** Both setting Ω_*EC*50_ to zero or omitting Ω_*EC*50_ entirely leads to the same values for all components (only Ω_*EC*50_ = 0 shown here). **Sc5** Ω_*EC*50_ is removed and the interindividual variability is re-introduced by “fixing the corresponding *P*_*EC*50_s to covariates”, see 2.3. Here, all terms containing contributions from Ω_*EC*50_ change significantly.

## 5 Discussion

In this paper, we analyzed the behavior of the Objective Function Value (OFV) when parameters are fixed to specific values and the remaining parameters are refitted. We identified a distinct behavior for the population approach compared to the non-population approach.

Firstly, we found that due to the explicit dependency of the OFV on population parameters, fixing some to certain values and refitting the rest, gives rise to divergent terms within the OFV. We demonstrated that these divergent terms cancel out, leading to a continuous behavior of the overall OFV and by this ensuring that a necessary condition for the LRT is met.

Secondly, we explored the case of fixing individual parameters (*P* s) and observed an unexpected behavior. Typically, it is anticipated that following a fit, the OFV remains the same or increases when a subset of parameters is fixed and other parameters are refitted. We confirmed that this is true for the non-population approach and also for population parameters in the population approach.

However, the OFV decreases when fixing individual parameters within the population approach to the estimated individual parameters (EBEs), although the population and individual predictions are identical. We conclude that valid model comparisons cannot be made when individual parameters are fixed.

### What does fixing of individual parameters imply?

In the following we discuss possibilities to introduce certain values for individual parameters (*P* s) and an alternative way to look at fixing individual parameters.

Observe the case when the baseline of an observed variable is included in the model as a parameter, the baseline has been measured for each individual and should be used: it is not recommended to fix the individual parameters (*P* s) to the measured individual baselines. Instead, it is advisable to introduce the measured baselines as covariates. A thorough analysis of the appropriate procedure can be found in [4].

For a more general approach, consider the case where individual parameters, *P*_*i*_s, are to be fixed to a set of values with variability Ω_*i*_. Fixing *P*_*i*_s, e.g. to previously fitted values *P*_*est,i*_ (EBEs), resembles the introduction of covariates with variability Ω_*i*_. Due to this similarity, an exploratory analysis comparing different scenarios was conducted:

- Fit the model (original fit)
- Replace the *P* s to the previously estimated individual parameters *P*_*est*_ (EBEs) and remove the corresponding Ω (same as Sc5 in 2.4)
- Introduce the *P*_*est*_s (EBEs) as covariates
- Add increasing levels of Gaussian noise to the *P*_*est*_s (EBEs) and use these as covariates

In the first three scenarios, the individual and population prediction values, and hence the fit to the data (Terms 1 and 2), remains unchanged. Interestingly, even though the second and third scenarios give identical individual and population predictions, the OFV shows a significant improvement compared to the OFV of the original fit. The second scenario is discussed in this work in some detail. In the third scenario, Ω is estimated to be very small, as the variability is explained by the introduction of the covariates. In the last scenario, where the covariates gradually deviate from the *P*_*est*_s, the OFV worsens as expected.

As discussed in this work, it is counterintuitive to observe a significant improvement of the OFV when the parameters (*P* s) are fixed to the *P*_*est*_s of the original fit, since the fit to the data stays the same. A tentative conclusion is that fixing individual parameters *P* s in the population approach cannot be compared to fixing classical (population) parameters. Instead, it should be considered a modification to the model where the external variability is explained by covariates, resulting in a better model. The fact that introducing covariates explains a population parameter (the corresponding variability Ω) and thus the model has a parameter less, suggests a change by one degree of freedom.

### Outlook

The exact implications of introducing covariates and the following change in the degree of freedom merit further exploration.

Further investigations could lead to a generalization of this work. Firstly, although the proof is currently made for a diagonal Ω matrix, we assume it would work analogously for a full matrix, but this remains to be proven. Secondly, we have analyzed the behaviour for FOCE (and in the numerical example: FOCE-I) with an additional error, but believe the theoretical analysis to be analogous for FOCE-I and the conclusions to be true for FOCE-I and different error models. This would need to be shown. Thirdly, we have only examined a necessary condition for a test statistic to be applied, i.e., ensuring the test statistic is continuous, but the sufficiency of these conditions requires further analysis.

### Summary

In summary, we have proven that a necessary condition for valid model comparisons is satisfied when fixing population parameters, as the change in likelihood is continuous. Fixing individual parameters (*P* s) on the other hand can lead to an unexpected improvement of the likelihood which would impede a valid test statistic. A viable approach to modifying individual parameters is the introduction of covariates (see for example, [3]). When covariates are introduced, the improvement in the OFV does not necessarily reflect an improved fit to the data but indicates that external variability, estimated by the corresponding population parameter Ω, is explained by the covariates. The change in the degree of freedom due to the introduction of covariates warrants further investigation.

## Acknowledgments

The authors wish to thank Philippe Pierrillas, Anna Fochesato, Antoine Soubret, Nicolas Frey and Hanna Silber Baumann (Pharmaceutical Sciences, Roche Innovation Center, Basel) for their insightful discussions and critical reading of the manuscript, which greatly improved its quality.

## Author Contributions

L.M.A. conducted the analysis with critical guidance and input from B.S.; both authors jointly wrote the manuscript.

