## Supplementary Code and Figures for "Theoretical and numerical investigation of the consistency of model comparisons in pharmacometrics"

### A Overview of key logarithmic and determinant calculation techniques applied in this work

- The log of a product is the addition of the elements where the logarithm is taken of

$$\log(a \cdot b) = \log(a) + \log(b) \tag{22}$$

- The determinant of a diagonal matrix, $A$ is the product of its elements

$$|A| = \left| \begin{bmatrix} a_1 & 0 & ... & 0 \\ 0 & a_2 & ... & 0 \\ ... & ... & ... & ... \\ 0 & 0 & ... & a_n \end{bmatrix} \right| = a_1 \cdot a_2 \cdot ... \cdot a_n \tag{23}$$

- The determinant of any 2x2 matrix is

$$\left| \begin{bmatrix} a & b \\ c & d \end{bmatrix} \right| = ad - bc \tag{24}$$

- The determinant of a $nxn$ matrix $A$ is calculated as follows. Let $minor(A)_{ij}$ be the determinant of the matrix which results from deleting the $i$th row and the $j$th column of matrix $A$. Let further $C_{ij}$ be the cofactor of matrix $A$, where

$$C_{ij} = (-1)^{i+j} minor(A)_{ij}. \tag{25}$$

Then the determinant of a $nxn$ with $n \geq 2$ is defined as

$$|A| = \sum_{j=1}^{n} a_{ij} C_{ij} = \sum_{i=1}^{n} a_{ij} C_{ij} \tag{26}$$

where the first expression expands the determinant along any chosen $i$th row and the second expression expands the determinant across any chosen $j$th column.

Example: We calculate the determinant of the $3x3$ matrix

$$A = \begin{bmatrix} 1 & 2 & 3 \\ 4 & 3 & 2 \\ 3 & 2 & 1 \end{bmatrix} \tag{27}$$

.

To this end we expand along the first row:

$$|A| = 1 \cdot (-1)^{1+1} \left| \begin{bmatrix} 3 & 2 \\ 2 & 1 \end{bmatrix} \right| + 2 \cdot (-1)^{1+2} \left| \begin{bmatrix} 4 & 2 \\ 3 & 1 \end{bmatrix} \right| + 3 \cdot (-1)^{1+3} \left| \begin{bmatrix} 4 & 3 \\ 3 & 2 \end{bmatrix} \right| \tag{28}$$

and applying the determinant rule for $2x2$ matrix we obtain

$$|A| = 1 \cdot (-1)^{1+1} \cdot (3-4) + 2 \cdot (-1)^{1+2} \cdot (4-6) + 3 \cdot (-1)^{1+3} \cdot (8-9) = -1 + 4 - 3 = 0 \tag{29}$$

### B Model for numerical example

The 2CMT model with absorption is fixed throughout the whole simulation and fitting and takes the form

$$\frac{dA_1}{dt} = \text{BOLUS} - K_A \cdot A_0$$
$$\frac{dA_2}{dt} = K_A \cdot A_0 - K_{12} * A_1 + K_{21} * A_2 - K_E * A_1$$
$$\frac{dA_3}{dt} = K_{12} * A_1 - K_{21} * A_2$$
$$C_{Drug} = \frac{A_2}{V_1}$$

with

$$KE = \frac{CL}{V1}$$
$$K23 = \frac{Q}{V1}$$
$$K32 = \frac{Q}{V2}$$

.

where $A_1$, $A_2$ and $A_3$ are the amounts of drug in the respective compartments and $C_{Drug}$ is the concentration in the central compartment. The indirect response model is given as

$$\frac{d\text{Response}}{\text{dt}} = \text{k}_{in} - \left(1 + \frac{\text{E}_{\text{max}} \cdot \text{C}_{\text{Drug}}}{\text{EC50} + \text{C}_{\text{Drug}}}\right) \text{K}_{out} \cdot \text{Response} \tag{30}$$

.

The following parameters were used for simulation:

| Parameter | value [a.u.] |
|---|---|
| $K_A$ | 1 |
| $CL$ | 1 |
| $V_1$ | 10 |
| $Q$ | 1 |
| $V_2$ | 10 |
| $K_{out}$ | 0.01 |
| $Response_{BL}$ | 10 |
| $IMAX$ | 1 |
| $EC50$ | 2.5 |
| $\Omega_{Kout}$ | 0.01 |
| $\Omega_{ResponseBL}$ | 0.05 |
| $\Omega_{EC50}$ | 0.1 |
| Residual variability | |
| additional error | 0.1 |

Table 2: Caption

The script for simulating the data can be found in C.1.

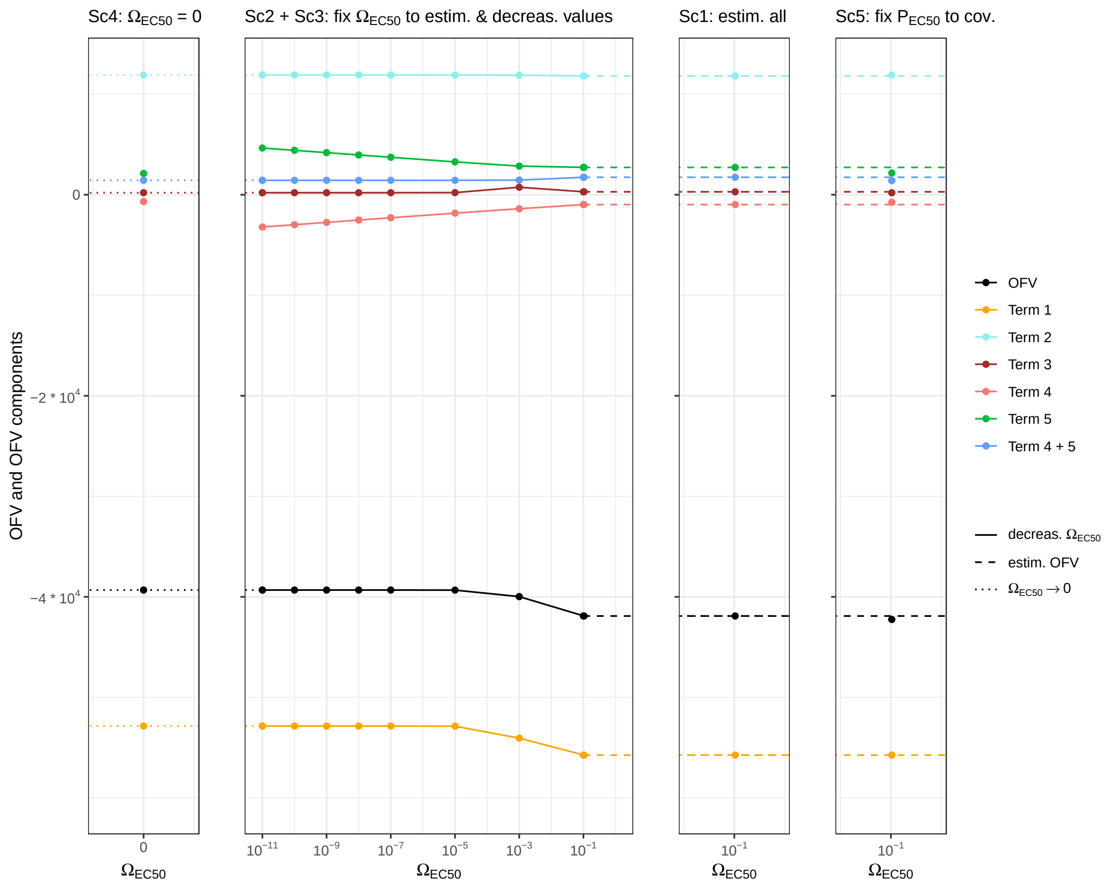


Figure 4: Values of OFV components for different fitting scenarios (Sc1-Sc5) where $\Omega_{EC50}$ is estimated, fixed or omitted. **Sc1** all parameters are estimated simultaneously. **Sc2** $\Omega_{EC50}$ and $\Theta_{EC50}$ are fixed to the estimated value from Sc1 and all other parameters are refitted. **Sc3** $\Omega_{EC50}$ is fixed to decreasing values and a contiuous change of components can be seen. A divergence of term 4 and 5 can be seen. **Sc4** Both setting $\Omega_{EC50}$ to zero or omitting $\Omega_{EC50}$ entirely leads to the same values for all components (only $\Omega_{EC50} = 0$ shown here). **Sc5** $\Omega_{EC50}$ is removed and the interindividual variability is re-introduced by "fixing the corresponding $P_{EC50}$s to covariates", see 2.3. Here, all terms containing contributions from $\Omega_{EC50}$ change significantly.

### C NONMEM and R scripts for the different scenarios and their analysis

#### C.1 Script to produce the simulated data set

```
## Simulate data for a two cmt model ###################################

# needed libraries

library(rxode2)
library(dplyr)
library(readr)
library(ggplot2)



#### make own Cov_df ##################################################################


uniqueIDs<-seq(1,100)
Nsubj<-length(uniqueIDs)



########## Model #####################################################################################


mod1 <-rxode2({

  # same PK parameters for all individuals
  KA=1
  CL=1
  V1= 10
  Q=1
  V2=10

  KE=CL/V1
  K23=Q/V1
  K32=Q/V2

  # indirect response parameters
  KOUT=ThetaKOUT*exp(eta.KOUT)
  RESPONSE_BL=ThetaRESPONSE_BL*exp(eta.RESPONSE_BL)
  IMAX=ThetaImax
  EC50=ThetaEC50*exp(eta.EC50)

  KIN   = KOUT * RESPONSE_BL
  C2 = A2/V1


  A1(0) = 0
  A2(0) = 0
  A3(0) = 0
  A4(0) = RESPONSE_BL



  d/dt(A1)   = - KA*A1
  d/dt(A2)   =   KA*A1 - K23*A2 + K32*A3 - KE*A2
  d/dt(A3)   =   K23*A2 - K32*A3
  d/dt(A4)  =    KIN - KOUT*A4*(1+(IMAX*(C2)/(EC50+C2)))

```

```
  A4_obs = A4 + ThetaAdd*A4.err

})



#####

sigma <- lotri::lotri(A4.err ~ 1)

theta <- c(ThetaKOUT    = 0.01,
           ThetaRESPONSE_BL    = 10,
           ThetaImax    = 1,
           ThetaEC50   =  2.5,
           ThetaAdd   =  0.1
)


####### Covariance matrix for IIV
omega <- lotri::lotri(eta.KOUT  ~  0.01,
                      eta.RESPONSE_BL  ~ 0.05,
                      eta.EC50  ~  0.1)

##### simple  Simulation
    ######################################################################


# one dose
dose_times_lean<-c(0)
# determine sampling pattern
Sampling_Pk<-c(seq(0,24,2),seq(26,240,2))


ev_tb  <- eventTable()%>%
  et(dose=100, time=0)%>%
  et(id=uniqueIDs)%>%
  add.sampling(Sampling_Pk)%>%
  as_tibble()



seed1 <- 12345
PD_data <- rxSolve(mod1, ev_tb, theta, omega=omega, sigma=sigma, seed=seed1
    )

# check output with plot
ggplot(PD_data,aes(x=time,y=(A4_obs),group=id))+geom_line()

##### Make NONMEM data set from simulated data set
    ###########################

PD_data_A1<-PD_data[,c("id","time","A4_obs")]
names(PD_data_A1)<-c("ID","TIME","DV")
PD_data_A1$CMT<-2
PD_data_A1$AMT<-0
PD_data_A1$EVID<-0
PD_data_A1<-cbind(PD_data_A1,PD_data[,c("KOUT","RESPONSE_BL","IMAX","EC50")
    ])


# create table with  dosing events and combine with NONMEM data set

tab_unique<-do.call(rbind,lapply(unique(PD_data_A1$ID),function(id){
  return(subset(PD_data_A1,ID==id)[1,])}))

```

```
PD_data_dose=data.frame(ID=seq(1,100),TIME=0,DV=0,CMT=1,AMT=100,EVID=1)
PD_data_dose<-cbind(PD_data_dose,tab_unique[,c("KOUT","RESPONSE_BL","IMAX","EC50")])

PD_data<-rbind(PD_data_A1,PD_data_dose)
PD_data <- PD_data %>% arrange(CMT) %>% arrange(TIME) %>% arrange(ID)

# add a column with the baseline value of the response
PD_data_new<-list()
for ( i in 1:100){
  PD_data_i<-subset(PD_data,ID==i)
  BL<-subset(subset(PD_data_i,TIME==0),CMT==2)$DV
  PD_data_i$RESPONSE_BL=BL
  PD_data_new[[i]]<-PD_data_i
}
PD_data_new<-do.call(rbind,PD_data_new)

# flag baseline values
PD_data_new$BLFLAG<-0
PD_data_new$BLFLAG[which(PD_data_new$TIME==0 & PD_data_new$CMT==2)]<-1

ggplot(PD_data_new,aes(x=TIME,y=DV))+geom_point()

write.csv(PD_data_new,"Simulated_PD_data.csv",sep = ",", row.names = FALSE, na = "", quote=FALSE)
```

To the simulated data set, we add a column with the estimated individual parameters for EC50 (EBEs) from Scenario 1 for which NONMEM was used (see C.2). These EBEs can subsequently be used to fix individual parameters $P_{EC50,i}$, in Scenario 5, as described in 2.3.

```


# take result from NONMEM run
data = as.data.frame(as.matrix(read.table("run1.fit", skip=1, header=TRUE)))

# take simulated data set
tab<-read.csv("Simulated_PD_data.csv")

# add matching individual EC50 value to each individual
tab_new<-do.call(rbind,lapply(unique(tab$ID),function(id){
  subdata<-subset(data,ID==id)
  subtab<-subset(tab,ID==id)
  subtab$IC50est<-subdata$EC50[1]
  return(subtab)
}))

write.table(tab_new,"Simulated_PD_data_estEC50step2.csv",sep = ",", row.names = FALSE, na = "", quote=FALSE)
```

#### C.2 NONMEM scripts for the different scenarios

##### C.2.1 Scenario 1

```
$PROBLEM   PKPD
$INPUT     ID TIME DV CMT AMT EVID KOUT_Ind RESPONSE_BL_Ind IMAX_Ind EC50_Ind



$DATA    Simulated_PD_data.csv
IGNORE=@



```

```
$SUBROUTINE ADVAN13 TOL=12
$MODEL   COMP=(SC) ;
         COMP=(RESPONSE) ;
     COMP=(PLASMA) ;
     COMP=(PERIPH) ;


$PK

KA  = 1
CL  = 1
V1  = 10
Q   = 1
V2  = 10

KE=CL/V1
K23=Q/V1
K32=Q/V2

KOUT=THETA(1)*EXP(ETA(1))
RESPONSE_BL=THETA(2)*EXP(ETA(2))
IMAX=THETA(3)
EC50=THETA(4)*EXP(ETA(3))

KIN   = KOUT * RESPONSE_BL

A_0(1)  = 0
A_0(2)  = RESPONSE_BL
A_0(3)  = 0
A_0(4)  = 0


$DES
C2  = A(3)/V1            ;

DADT(1)  = - KA*A(1)
DADT(2)  =   KIN - KOUT*(1+IMAX*C2/(EC50+C2))*A(2)
DADT(3)  =   KA*A(1) - K23*A(3) + K32*A(4) - KE*A(3)
DADT(4)  =   K23*A(3) - K32*A(4)


$ERROR

RESPONSE = A(2)

ADD_RESPONSE = THETA(5);



IPRED  = RESPONSE
W      = ADD_RESPONSE ; additive
Y      = IPRED + W*ERR(1)


IRES=IPRED-DV
IWRES = IRES/W


$THETA

(0,0.011)    ;KOUT
```

```
(0,9)      ;RESPONSE_BL
(0,1.1)      ;IMAX
(0,3)        ;EC50
(0, 0.012)     ; additive error

$OMEGA
0.011      ; KOUT
0.051      ; RESPONSE_BL
0.11       ; EC50




$SIGMA
1 FIX

$EST  MAXEVAL=9999 NSIG=3 SIGL=9 PRINT=5 METHOD=1 INTER NOABORT
NOTHETABOUNDTEST NOOMEGABOUNDTEST NOSIGMABOUNDTEST
MSFO = run01.msfo
$COV MATRIX=S
$TABLE ID TIME DV CMT AMT EVID  PRED IPRED IRES IWRES NPDE CWRES
    KOUT KOUT_Ind  RESPONSE_BL_Ind RESPONSE_BL  EC50 EC50_Ind  G11
    G21 G31 H11 ETA1 ETA2 ETA3
NOPRINT ONEHEADER NOAPPEND
FILE = run1.fit
```

##### C.2.2 Scenario 2

```
$PROBLEM   PKPD
$INPUT     ID TIME DV CMT AMT EVID KOUT_Ind RESPONSE_BL_Ind IMAX_
    Ind EC50_Ind



$DATA    Simulated_PD_data.csv
IGNORE=@



$SUBROUTINE ADVAN13 TOL=12
$MODEL   COMP=(SC) ;
         COMP=(RESPONSE) ;
     COMP=(PLASMA) ;
     COMP=(PERIPH) ;




$PK

KA = 1
CL = 1
V1 = 10
Q  = 1
V2 = 10

KE=CL/V1
K23=Q/V1
K32=Q/V2

```

```
KOUT=THETA(1)*EXP(ETA(1))
RESPONSE_BL=THETA(2)*EXP(ETA(2))
IMAX=THETA(3)
EC50=THETA(4)*EXP(ETA(3))

KIN  = KOUT * RESPONSE_BL

A_0(1) = 0
A_0(2) = RESPONSE_BL
A_0(3) = 0
A_0(4) = 0


$DES
C2 = A(3)/V1            ;  mcg/l oder ng/ml


DADT(1) = - KA*A(1)
DADT(2) =   KIN - KOUT*(1+IMAX*C2/(EC50+C2))*A(2)
DADT(3) =   KA*A(1) - K23*A(3) + K32*A(4) - KE*A(3)
DADT(4) =   K23*A(3) - K32*A(4)


$ERROR

RESPONSE = A(2)

ADD_RESPONSE = THETA(5);



IPRED = RESPONSE
W     = ADD_RESPONSE
Y     = IPRED + W*ERR(1)


IRES=IPRED-DV
IWRES = IRES/W


$THETA
(0,0.011)       ;KOUT
(0,9)     ;RESPONSE_BL
(0,1.1)     ;IMAX
(0,2.34)  FIX ;EC50
(0, 0.012)    ; additive error

$OMEGA
0.011     ; KOUT
0.051     ; RESPONSE_BL
0.104 FIX   ; EC50




$SIGMA
1 FIX

$EST  MAXEVAL=9999 NSIG=3 SIGL=9 PRINT=5 METHOD=1 INTER NOABORT
NOTHETABOUNDTEST NOOMEGABOUNDTEST NOSIGMABOUNDTEST
MSFO = run01.msfo
```

```
$COV MATRIX=S
$TABLE ID TIME DV CMT AMT EVID  PRED IPRED IRES IWRES NPDE CWRES
    KOUT KOUT_Ind  RESPONSE_BL_Ind RESPONSE_BL  EC50 EC50_Ind  G11
    G21 G31 H11 ETA1 ETA2 ETA3
NOPRINT ONEHEADER NOAPPEND
FILE = run1.fit
```

##### C.2.3 Scenario 3

```
$PROBLEM   PKPD
$INPUT      ID TIME DV CMT AMT EVID KOUT_Ind RESPONSE_BL_Ind IMAX_
    Ind EC50_Ind



$DATA    Simulated_PD_data.csv
IGNORE=@



$SUBROUTINE ADVAN13 TOL=12
$MODEL   COMP=(SC) ;
         COMP=(RESPONSE) ;
     COMP=(PLASMA) ;
     COMP=(PERIPH) ;



$PK

KA  = 1
CL  = 1
V1  = 10
Q   = 1
V2  = 10

KE=CL/V1
K23=Q/V1
K32=Q/V2

KOUT=THETA(1)*EXP(ETA(1))
RESPONSE_BL=THETA(2)*EXP(ETA(2))
IMAX=THETA(3)
EC50=THETA(4)*EXP(ETA(3))

KIN   = KOUT * RESPONSE_BL

A_0(1)  = 0
A_0(2)  = RESPONSE_BL
A_0(3)  = 0
A_0(4)  = 0


$DES
C2  = A(3)/V1             ;


DADT(1)  = - KA*A(1)
DADT(2)  =   KIN - KOUT*(1+IMAX*C2/(EC50+C2))*A(2)
DADT(3)  =   KA*A(1) - K23*A(3) + K32*A(4) - KE*A(3)
```

```
DADT(4) =   K23*A(3) - K32*A(4)



$ERROR

RESPONSE = A(2)

ADD_RESPONSE = THETA(5);



IPRED = RESPONSE

W      = ADD_RESPONSE ; additive
Y      = IPRED + W*ERR(1)


IRES=IPRED-DV
IWRES = IRES/W


$THETA
(0,0.011)    ;KOUT
(0,9)      ;RESPONSE_BL
(0,1.1)      ;IMAX
(0,1.1)      ;EC50
(0, 0.012)     ;additive error

$OMEGA
0.011      ; KOUT
0.051      ; RESPONSE_BL
0.001   FIX    ; EC50  (or smaller values up to 1e-11)




$SIGMA
1 FIX

$EST  MAXEVAL=9999 NSIG=3 SIGL=9 PRINT=5 METHOD=1 INTER NOABORT
NOTHETABOUNDTEST NOOMEGABOUNDTEST NOSIGMABOUNDTEST
MSFO = run01.msfo
$COV MATRIX=S
$TABLE ID TIME DV CMT AMT EVID  PRED IPRED IRES IWRES NPDE CWRES
    KOUT KOUT_Ind  RESPONSE_BL_Ind RESPONSE_BL  EC50 EC50_Ind  G11
    G21 G31 H11 ETA1 ETA2 ETA3
NOPRINT ONEHEADER NOAPPEND
FILE = run1.fit
```

##### C.2.4 Scenario 4

```
$PROBLEM   PKPD Omega set to zero
$INPUT     ID TIME DV CMT AMT EVID KOUT_Ind RESPONSE_BL_Ind IMAX_
    Ind EC50_Ind



$DATA    Simulated_PD_data.csv
IGNORE=@
```

```
$SUBROUTINE ADVAN13 TOL=12
$MODEL  COMP=(SC) ;
        COMP=(RESPONSE) ;
    COMP=(PLASMA) ;
    COMP=(PERIPH) ;


$PK

KA  = 1
CL  = 1
V1  = 10
Q   = 1
V2  = 10

KE=CL/V1
K23=Q/V1
K32=Q/V2

KOUT=THETA(1)*EXP(ETA(1))
RESPONSE_BL=THETA(2)*EXP(ETA(2))
IMAX=THETA(3)
EC50=THETA(4)*EXP(ETA(3))

KIN   = KOUT * RESPONSE_BL

A_0(1)  = 0
A_0(2)  = RESPONSE_BL
A_0(3)  = 0
A_0(4)  = 0


$DES
C2  = A(3)/V1


DADT(1)  = - KA*A(1)
DADT(2)  =   KIN - KOUT*(1+IMAX*C2/(EC50+C2))*A(2)
DADT(3)  =   KA*A(1) - K23*A(3) + K32*A(4) - KE*A(3)
DADT(4)  =   K23*A(3) - K32*A(4)


$ERROR

RESPONSE = A(2)

ADD_RESPONSE = THETA(5);



IPRED = RESPONSE
W     = ADD_RESPONSE ; additive
Y     = IPRED + W*ERR(1)


IRES=IPRED-DV
IWRES = IRES/W
```

```

$THETA
(0,0.011)    ;KOUT
(0,9)      ;RESPONSE_BL
(0,1.1)      ;IMAX
(0,1.1)      ;EC50
(0, 0.012)     ; add error

$OMEGA
0.011      ; KOUT
0.051      ; RESPONSE_BL
0 FIX        ; EC50




$SIGMA
1 FIX

$EST  MAXEVAL=9999 NSIG=3 SIGL=9 PRINT=5 METHOD=1 INTER NOABORT
NOTHETABOUNDTEST NOOMEGABOUNDTEST NOSIGMABOUNDTEST
MSFO = run01.msfo
$COV MATRIX=S
$TABLE ID TIME DV CMT AMT EVID  PRED IPRED IRES IWRES NPDE CWRES KOUT KOUT_Ind  RESPONSE_BL_Ind RESPONSE_BL  EC50 EC50_Ind  G11 G21 G31 H11 ETA1 ETA2 ETA3
NOPRINT ONEHEADER NOAPPEND
FILE = run1.fit





; Second script for Omitted Omega


$PROBLEM  PKPD Omega omitted
$INPUT     ID TIME DV CMT AMT EVID KOUT_Ind RESPONSE_BL_Ind IMAX_Ind EC50_Ind



$DATA    Simulated_PD_data.csv
IGNORE=@



$SUBROUTINE ADVAN13 TOL=12
$MODEL  COMP=(SC) ;
        COMP=(RESPONSE) ;
    COMP=(PLASMA) ;
    COMP=(PERIPH) ;


$PK

KA = 1
CL = 1
V1 = 10
Q  = 1
V2 = 10
```

```
KE=CL/V1
K23=Q/V1
K32=Q/V2

KOUT=THETA(1)*EXP(ETA(1))
RESPONSE_BL=THETA(2)*EXP(ETA(2))
IMAX=THETA(3)
EC50=THETA(4)

KIN  = KOUT * RESPONSE_BL

A_0(1) = 0
A_0(2) = RESPONSE_BL
A_0(3) = 0
A_0(4) = 0


$DES
C2 = A(3)/V1


DADT(1) = - KA*A(1)
DADT(2) =   KIN - KOUT*(1+IMAX*C2/(EC50+C2))*A(2)
DADT(3) =   KA*A(1) - K23*A(3) + K32*A(4) - KE*A(3)
DADT(4) =   K23*A(3) - K32*A(4)



$ERROR

RESPONSE = A(2)

ADD_RESPONSE = THETA(5);



IPRED = RESPONSE

W      = ADD_RESPONSE ; additive
Y      = IPRED + W*ERR(1)


IRES=IPRED-DV
IWRES = IRES/W


$THETA
(0,0.011)    ;KOUT
(0,9)      ;RESPONSE_BL
(0,1.1)      ;IMAX
(0,1.1)      ;EC50
(0, 0.012)     ; add error

$OMEGA
0.011      ; KOUT
0.051      ; RESPONSE_BL
```

```

$SIGMA
1 FIX

$EST  MAXEVAL=9999 NSIG=3 SIGL=9 PRINT=5 METHOD=1 INTER NOABORT
NOTHETABOUNDTEST NOOMEGABOUNDTEST NOSIGMABOUNDTEST
MSFO = run01.msfo
$COV MATRIX=S
$TABLE ID TIME DV CMT AMT EVID  PRED IPRED IRES IWRES NPDE CWRES KOUT KOUT_Ind  RESPONSE_BL_Ind RESPONSE_BL  EC50 EC50_Ind  G11 G21  H11 ETA1 ETA2
NOPRINT ONEHEADER NOAPPEND
FILE = run1.fit
```

##### C.2.5 Scenario 5

```
$PROBLEM  PLASMA PKPD
$INPUT     ID TIME DV CMT AMT EVID KOUT_Ind RESPONSE_BL_Ind IMAX_Ind EC50_Ind RESPONSE_BL BLFLAG EC50est

$DATA    Simulated_PD_data_estEC50step2.csv
IGNORE=@

$SUBROUTINE ADVAN13 TOL=12
$MODEL  COMP=(SC) ;
        COMP=(RESPONSE) ;
    COMP=(PLASMA) ;
    COMP=(PERIPH) ;

$PK

KA  = 1
CL  = 1
V1  = 10
Q   = 1
V2  = 10

KE=CL/V1
K23=Q/V1
K32=Q/V2

KOUT=THETA(1)*EXP(ETA(1))
RESPONSE_BL=THETA(2)*EXP(ETA(2))
IMAX=THETA(3)
EC50=EC50est

KIN   = KOUT * RESPONSE_BL

A_0(1)  = 0
A_0(2)  = RESPONSE_BL
A_0(3)  = 0
A_0(4)  = 0

```

```
$DES
C2 = A(3)/V1


DADT(1) = - KA*A(1)
DADT(2) =   KIN - KOUT*(1+IMAX*C2/(EC50+C2))*A(2)
DADT(3) =   KA*A(1) - K23*A(3) + K32*A(4) - KE*A(3)
DADT(4) =   K23*A(3) - K32*A(4)



$ERROR

RESPONSE = A(2)

ADD_RESPONSE = THETA(4);

IPRED = RESPONSE

W      = ADD_RESPONSE ; additive
Y      = IPRED + W*ERR(1)


IRES=IPRED-DV
IWRES = IRES/W


$THETA

9.92605E-03 FIX      ;KOUT
1.01543E+01 FIX      ;RESPONSE_BL
9.81382E-01 FIX      ;IMAX

9.92710E-02              ; add error

$OMEGA
8.11712E-03 FIX      ; KOUT
4.67742E-02 FIX      ; RESPONSE_BL





$SIGMA
1 FIX

$EST  MAXEVAL=0 NSIG=3 SIGL=9 PRINT=5 METHOD=1 INTER NOABORT
NOTHETABOUNDTEST NOOMEGABOUNDTEST NOSIGMABOUNDTEST
MSFO = run01.msfo
$COV MATRIX=S
$TABLE ID TIME DV CMT AMT EVID  PRED IPRED IRES IWRES NPDE CWRES
    KOUT KOUT_Ind  RESPONSE_BL_Ind RESPONSE_BL  EC50 EC50_Ind  G11
    G21  H11 ETA1 ETA2
NOPRINT ONEHEADER NOAPPEND
FILE = run1.fit
```

#### C.3 Scripts for the calculation of OFV components

Scripts for calculating OFV components from NONMEM output. The first script is detailing the calculation for a 3x3 $\Omega$ matrix using functions defined in the second script - both are adapted from

[1].

```r
library(dplyr)
source("BAE_YIM_functions.R")

# Control file name without extension
CtlName = "Run1"

# read in NONMEM results
DATA = as.data.frame(as.matrix(read.table("run1.fit", skip=1, header=TRUE)))
DATA= as.matrix(DATA%>%filter(!EVID==1))

# read in population parameter output by NONMEM
EXT = as.matrix(read.table(paste0(CtlName, ".ext"), skip=1, header=TRUE))
last_pos_iteration<-max(which(EXT[,c("ITERATION")]>0))
# Omega matrix
OM = ltv2mat(EXT[last_pos_iteration, c("OMEGA.1.1.", "OMEGA.2.1.", "OMEGA.2.2.","OMEGA.3.1.", "OMEGA.3.2.","OMEGA.3.3.")])
# Sigma matrix
SG = matrix(EXT[last_pos_iteration, "SIGMA.1.1."], nrow=1, ncol=1)

# read in EBE output by NONMEM
EBE = as.matrix(read.table(paste0(CtlName, ".phi"), skip=1, header=TRUE))

IDs = unique(DATA[,"ID"])
nID = length(IDs)
Term4 = determinant(OM, logarithm=TRUE)$modulus[[1]]
invOM = solve(OM)
CWRES = vector()

OFV_list<-lapply (1:nID, function(i){
  cID = IDs[i]
  DATi = DATA[DATA[,"ID"]==cID, ]
  EBEi = EBE[EBE[,"ID"]==cID, c("ETA.1.", "ETA.2.","ETA.3.")]
  Yi = DATi[,"DV"]
  Fi = DATi[,"IPRED"]
  Gi = DATi[, c("G11", "G21","G31")]
  Hi = DATi[, c("H11")]
  Vi = diag(diag(Hi %*% SG %*% t(Hi)))
  invVi = solve(Vi)
  Term1 = determinant(Vi, logarithm=TRUE)$modulus[[1]] # log(det(Vi))
  Term2 = t(Yi - Fi)%*% invVi %*%(Yi - Fi)
  Term3 = t(EBEi) %*% invOM %*% EBEi
  # Term4 = determinant(OM, logarithm=TRUE)$modulus[[1]]
  Term5 = log(det(invOM + t(Gi) %*% invVi %*% Gi))
  foce = Term1 + Term2 + Term3 + Term4 + Term5


  Ci = Gi %*% OM %*% t(Gi) + diag(diag(Hi %*% SG %*% t(Hi)))
  ginv = solve(Ci)
  term2 =  sec<-t(Yi - Fi + Gi %*% EBEi)%*%ginv%*%(Yi - Fi + Gi %*% EBEi)
  #CWRESi = mat.sqrt.inv(Ci) %*% (Yi - Fi + Gi %*% EBEi)
  fo = log(det(Ci)) + term2#t(CWRESi) %*% CWRESi
  #CWRES = append(CWRES, CWRESi)

 OFV_df<-data.frame(ID=cID,ResErr=Term1,Res=Term2,ResEta=Term3,OmegaErr=Term4,SecondDer=Term5,FOCE=foce,FO=fo)

 return(OFV_df)
})
OFV_DF<-do.call(rbind,OFV_list)

OFV_DF_final<-apply(OFV_DF,2,sum)
names(OFV_DF_final)<-c("ID","ResErr","Res","ResEta","OmegaErr","SecondDer","FOCE","FO")
```

```
#alternative names: ResErr=Term1,Res=Term2,ResEta=Term3,OmegaErr=Term4,
    SecondDer=Term5

write.csv(OFV_DF_final,"OFV_DF_final.csv")
```

Second script, functions adapted from [1].

```


##### adapted from  BAE YIM publication #########################333

SqrtInvCov = function(M)
{
  EigenResult = eigen(as.matrix(M))
  EigenVector = EigenResult$vectors
  EigenValues = abs(EigenResult$values)
  return(EigenVector %*% diag(1/sqrt(EigenValues)) %*% t(EigenVector))
}


Mx = function(M, x)
{
  e = eigen(as.matrix(M))
  return(e$vectors %*% diag(abs(e$values)^x) %*% t(e$vectors))
}



mat2ltv = function(mat)
{
  return(mat[upper.tri(mat,diag=TRUE)])
}

ltv2mat = function(vec)
{
  LENGTH = length(vec)
  DIM = round((sqrt(8*LENGTH+1)-1)/2,0)
  if (DIM*(DIM+1)/2 != LENGTH) return(NULL)
  mat = matrix(nrow=DIM, ncol=DIM)
  for (m in 1:DIM) {
    for (n in 1:DIM) {
      k = max(m,n)
      l = min(m,n)
      p = k * (k-1) / 2 + l
      mat[m,n] = vec[p]
    }
  }
  return(mat)
}


mat2utv = function(mat)
{
  return(mat[lower.tri(mat,diag=TRUE)])
}


utv2mat = function(vec)
{
  LENGTH = length(vec)
  DIM = as.integer(round((sqrt(8*LENGTH+1)-1)/2,0))
  if (DIM*(DIM+1)/2 != LENGTH) return(NULL)
  mat = matrix(nrow=DIM, ncol=DIM)
  for (m in 1:DIM) {
    for (n in 1:DIM) {
      k = min(m,n)
      l = max(m,n)
```

```
      p = (2*DIM - k + 2)*(k - 1)/2 + l - k + 1
      mat[m,n] = vec[p]
    }
  }
  return(mat)
}


ScaleVar = function(VarMat, dim1)
{
  M1 = chol(VarMat)
  V1 = diag(M1)
  M2 = abs(10 * (M1 - diag(V1, nrow=dim1))) + diag(V1/exp(0.1), nrow=dim1)
  return(t(M2))
}


DesclVar = function(mUCP, mSCL)
{
  nRow = dim(mUCP)[1]
  maT = matrix(nrow=nRow, ncol=nRow)

  for (i in 1:nRow) {
    for (j in 1:nRow) {
      if (i==j) {
        maT[i,j] = exp(mUCP[i,j]) * mSCL[i,j]
      } else if(i > j) {
        maT[i,j] = mUCP[i,j] * mSCL[i,j]
      } else {
        maT[i,j] = 0
      }
    }
  }
  return(maT %*% t(maT))
}


DECN = function(UCP)
{
  #Extern: nTheta, nEta, nEps
  #        ThetaL[nTheta]==LB, ThetaI[nTheta]==IE, ThetaU[nTheta]==UB
  #        alpha, OMscl, SGscl
  uTheta = UCP[1:e$nTheta]
  Theta = exp(uTheta - e$alpha)/(exp(uTheta - e$alpha) + 1)*(e$UB - e$LB) +
      e$LB
  uvOM  = UCP[(e$nTheta + 1):(e$nTheta + e$nEta*(e$nEta + 1)/2)]
  umOM  = utv2mat(uvOM)
  OM    = DesclVar(umOM, e$OMscl)
  ltvOM = mat2ltv(OM)

  udSG  = UCP[(e$nTheta + e$nEta*(e$nEta + 1)/2 + 1):(e$nTheta + e$nEta*(e$
    nEta + 1)/2 + e$nEps)]
  umSG  = diag(udSG, nrow=e$nEps)
  SG    = DesclVar(umSG, e$SGscl)
  dgSG  = diag(SG)
  return(c(Theta, ltvOM, dgSG))
}
```
